# Dispersal and evolution of the invasive snail *Pomacea canaliculata*, an intermediate host of *Angiostrongylus cantonensis*: A field study around its initial introduction site in China

**DOI:** 10.1101/2023.12.29.573677

**Authors:** Du Luo, Haitao Zhang, Dangen Gu, Xidong Mu, Hongmei Song, Yexin Yang, Meng Xu, Jianren Luo, Yinchang Hu

## Abstract

Eosinophilic meningitis caused by *Angiostrongylus cantonensis* is an emerging infectious disease in mainland China. The invasive snail (*Pomacea canaliculata*) is one of the main intermediate host of the zoonotic nematode. To gain insights into the spatial distribution, phenotypic variation and dispersal pattern of the snail populations. A field survey was conducted using modified nested sampling and selecting ditches as the main habitats. Snail distribution and abundance were displayed with predictions based on an inverse distance-weighted model. Correlation and spatial autocorrelation were analyzed among the measured parameters. The findings of this study demonstrate the well-established and abundant presence of *P. canaliculata* in ditches. A total of 564 sampling sites were assessed, with measurements taken on 10,145 snails having shell heights greater than 1.5 cm. The average population density was determined to be 20.31 ± 11.55 snails per square meter. Morphological analysis revealed an average snail body mass of 8.93 ± 3.95 g, shell height of 3.38 ± 0.66 cm, a sex ratio of 2.39 ± 1.01 female to male, and a shell color ratio of 9.34 ± 7.52 brown to yellow. Among these measurements, body mass was found to be significantly correlated with shell height (*r* = 0.88, *p* < 0.01) and shell color (*r* = 0.55, *p* < 0.05). Spatial-correlation analysis proved that shell height was the only factor significantly spatially autocorrelated (*MI* = 0.27, *z* = 2.20, *p* = 0.03), although weak autocorrelations appeared in body mass and shell color. The observed geographic variations of phenotypic traits indicated a human-mediated evolving process of the snail populations and a potential complexity of the parasite transmission system. These findings may enhance the assessment of the epidemiological health risk posed by angiostrongyliasis and inform strategies for controlling infectious snails.

## 1. Introduction

Invasive species and infectious disease are becoming more prevalent with increased globalization. Invasive disease vectors are spreading across geographical barriers as a result of human transport, land-use change, and climate change [1]. Vector-borne diseases, which are among the most common classes of emerging and re-emerging infectious diseases, represent a major threat to public health [2]. In particular, this freshwater snail is a vector of the parasitic nematode *Angiostrongylus cantonensis* (Chen, 1935), which commonly causes eosinophilic meningitis (angiostrongyliasis) in Southeast Asia, Australia, the Caribbean, South America and on Pacific Islands [3,4]. In the natural life cycle of the parasite *A. cantonensis*, snails serve as the primary intermediate hosts, while rats function as definitive hosts. The transmission of the parasite to human hinges on the pivotal role of intermediate hosts, particularly snails, in the infection process [5]. Angiostrongyliasis is recognized as an emerging infectious disease in China due to the invasive apple snail species serving as the primary source of infection [6]. A national survey confirmed that the predominant intermediate hosts of *A. cantonensis* were two invasive snails, *Achatina fulica* Bowdich, 1822 and *P. canaliculata* [7]. The latter species is more widely distributed and is well established in 16 provinces in mainland China. It was the cause of a series of angiostrongyliasis outbreaks, including the two largest involving 65 patients in Wenzhou in 1997 and 160 patients in Beijing in 2006. It appears that this invasive snail has facilitated the spread of the parasite and has led to the emergence of angiostrongyliasis.

The channeled apple snail *Pomacea canaliculata* (Lamarck, 1822) is a large aquatic gastropod mollusk in the family Ampullariidae. It is South American in origin and was introduced to Asian countries as a promising food crop in the early 1980s [8]. However, it soon escaped from aquaculture and established wild populations in many countries [9]. Also known as the golden apple snail, *P. canaliculata* is an extremely invasive molluskan pest and has established in many parts of the world including Asia, North America, islands of the Pacific, and Africa [10, 11]. This species was designated as one of the worst 100 globally invasive species in 2000 [12]. In 2003, *P. canaliculata* was also listed in the first list of 16 worst invasive species in China because of its devastating impact on agriculture, native species and human health. The snail’s extensive consumption of aquatic vegetation, accumulation of algal toxins and heavy metals, ability to spread widely and abundantly, and parasite burden have heightened the concerns surrounding its invasion [13].

*P. canaliculata* is a novel vector of *A. cantonensis* in China that was intentionally imported in 1981 for the purpose of aquaculture. The initial introduction site was Shaxi Town in Zhongshan City, Guangdong Province. Subsequently, the snails dispersed into various regions of south China as part of aquaculture promotion efforts involving single or multiple introductions [14]. The transmission process of *A. cantonensis* to humans typically involves the consumption of raw or undercooked intermediate hosts, such as infected snails, or inadvertent ingestion of contaminated vegetables or water containing infective larvae [15]. Although the snail is closely connected with outbreaks of eosinophilic meningitis, there has been no accurate monitoring of its natural abundance, distribution, and population changes since its introduction [16]. Phylogenetic evidence indicates that *Pomacea sp*. snails have been multiply and secondarily introduced intentionally into mainland China [17]. However, to date, there has been no detailed report on the natural distribution and dispersal pattern of these most invasive snails. Guangdong Province, the central area of the Pearl River Basin, bounded by the South China Sea, is the southern gateway of China. A provincial survey found that the highest rate of larval infection was 26.6%, and 86.4% investigated sites with infected snails, and that the prevalence of *A. cantonensis* in wild snails was a substantial risk for angiostrongyliasis in humans [18]. Studies on the susceptibility of *P. canaliculata* to *A. cantonensis* showed that infection rates were lowest in the yellow varieties and that earlier developmental stages of infected snails had a high mortality [19,20]. Temperature significantly affects infection intensity with an optimum temperature range of 20–30 °C and a lower critical temperature of 6.66 °C [21].

Many pest species exhibit huge fluctuations in population abundance. Even today, apple snails (*Pomacea sp*.) that pose a risk of spreading angiostrongyliasis are still being sold in the market with misleading food labeling [22]. Therefore, to improve response strategies to outbreaks and to develop effective control and management measures, it is necessary to understand the spatiotemporal population dynamics of such species [23]. Here, based on the previous research on change of phenotypic traits across different ecological scales in Guangdong province, we mainly report on a survey to explore the spatial variation of population density, morphology, sex ratio and shell color of the snails from ditch habitat [24]. The dispersal patterns, spatial distribution, and variation of phenotypic characteristics of the snail population were then analyzed and used for prediction. Recommendations are offered for control of population expansion and for preventing the further spread of angiostrongyliasis.

## 2. Materials and Methods

### 2.1. Design of the field survey

Zhongshan City, the initial site of introduction of the apple snail into mainland China, was established as the center of the study area. The survey field is shaped approximately in the form of a sector with a radius of 300 to 400 km. Geographically, the study area is located mainly in Guangdong Province and partially in Wuzhou City, Guangxi Province (Figure 1A). The survey was generally implemented between June and November in 2013 and 2014. Three sites in Zhaoqing, Guangzhou and Shaoguan City were repeatedly investigated from March to April during 2011 to 2014. Systematic random sampling was used in the present survey. In the Pearl River Delta, which was the inner area of the investigation, most of the cities were included as study sites to ensure a spatial resolution better than 50 × 50 km. In the outer area, the distance between each of the study sites was set to ≤ 100 km. A modified nested sampling method was implemented by randomly selecting three to five sample sites in each village, three villages in each town, and two or three towns in each city (Figure 1B). The geographical locations, distribution patterns of snails at each site, their abundance and phenotypes, as well as special environmental features were recorded.

**Figure 1.**
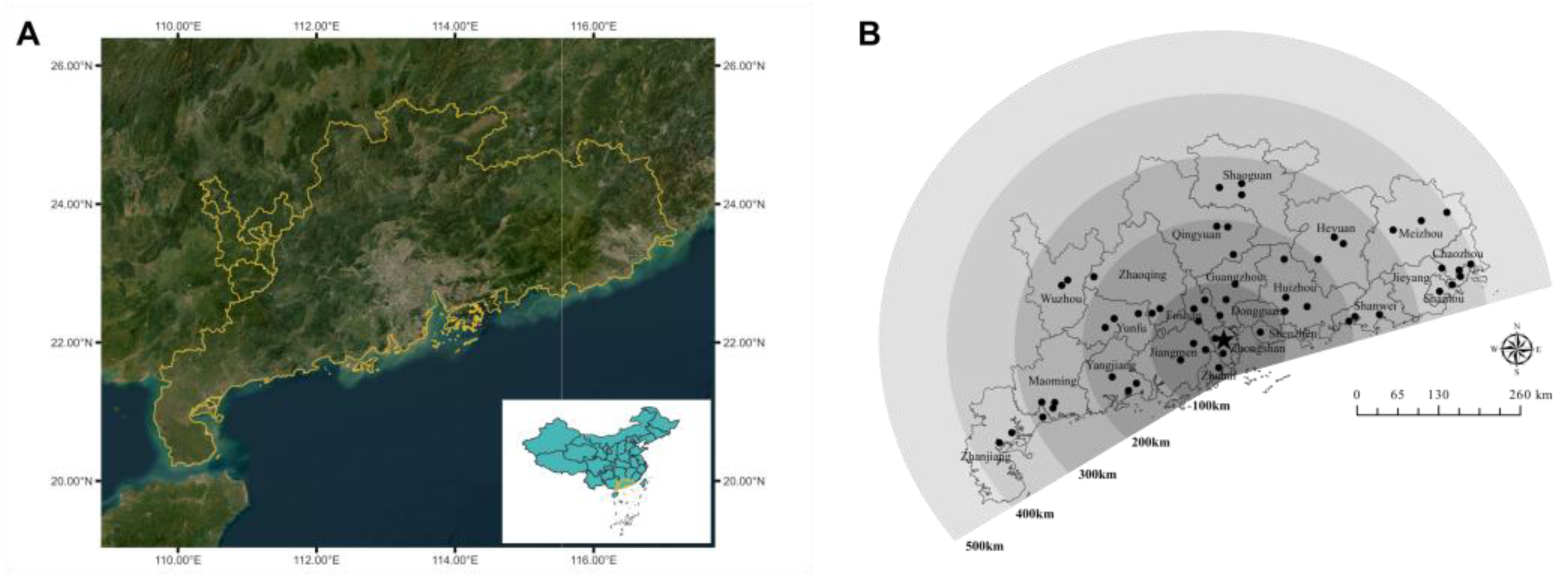
Sampling sites and study area in South China. This figure illustrates the location of the sampling sites within the study area. The research was conducted in South China (A), specifically in a sector (B) with a radius of approximately 400 km centered on Zhongshan city (marked by a star). Zhongshan city is the site of the initial introduction of *Pomacea canaliculata*, located on the coast of the South China Sea. Each point on the map represents a town where the investigation was carried out.

### 2.2. Population density and phenotypic variation of P. canaliculata

The habitats of *P. canaliculata* encompassed diverse environments, including rivers, streams, ditches, lakes, fish ponds, marshes, and rice paddies. A preliminary exploration of snail distribution and population density involved a thorough examination of these habitats. Ditches, functioning as waterways around rice fields and vegetable farms, were specifically scrutinized. Population densities and snail morphology in ditch habitats across various study sites were evaluated. During field investigations, a minimum of three sites were sampled within each sampling area. The sampling sites were confined to a 1 m² area, and the distance between each site ranged from 10 m to 30 m. Snails with a diameter exceeding approximately 1 cm were enumerated to calculate population density.

To reveal the distribution, population density, variation and dispersal patterns, a total of 564 sample sites from ditches were used to analyze population density. Snails with shell height > 1.5 cm from these sites were used to examine the morphological variation. Morphological indicators of body mass and shell height were measured according to Youens, et al. [25] and Hayes, et al. [26]. Shell surface, aperture morphology of the male individuals, operculum and body size were taken into account during observation. Shell color was classified as either banded-brown or yellow, identified by the color of the outer shell surface and the inner muscular foot [Figure 2]. The ratio of the two color forms is given as brown/yellow [27]. The sex ratio (female/male) of each sample was recorded for snails of shell height > 2.5 cm [28]. Associated aquatic snail species from the same habitat were also collected. Data for population density, body mass, shell height, sex ratio and color ratio are displayed geographically on the maps at city level.

**Figure 2.**
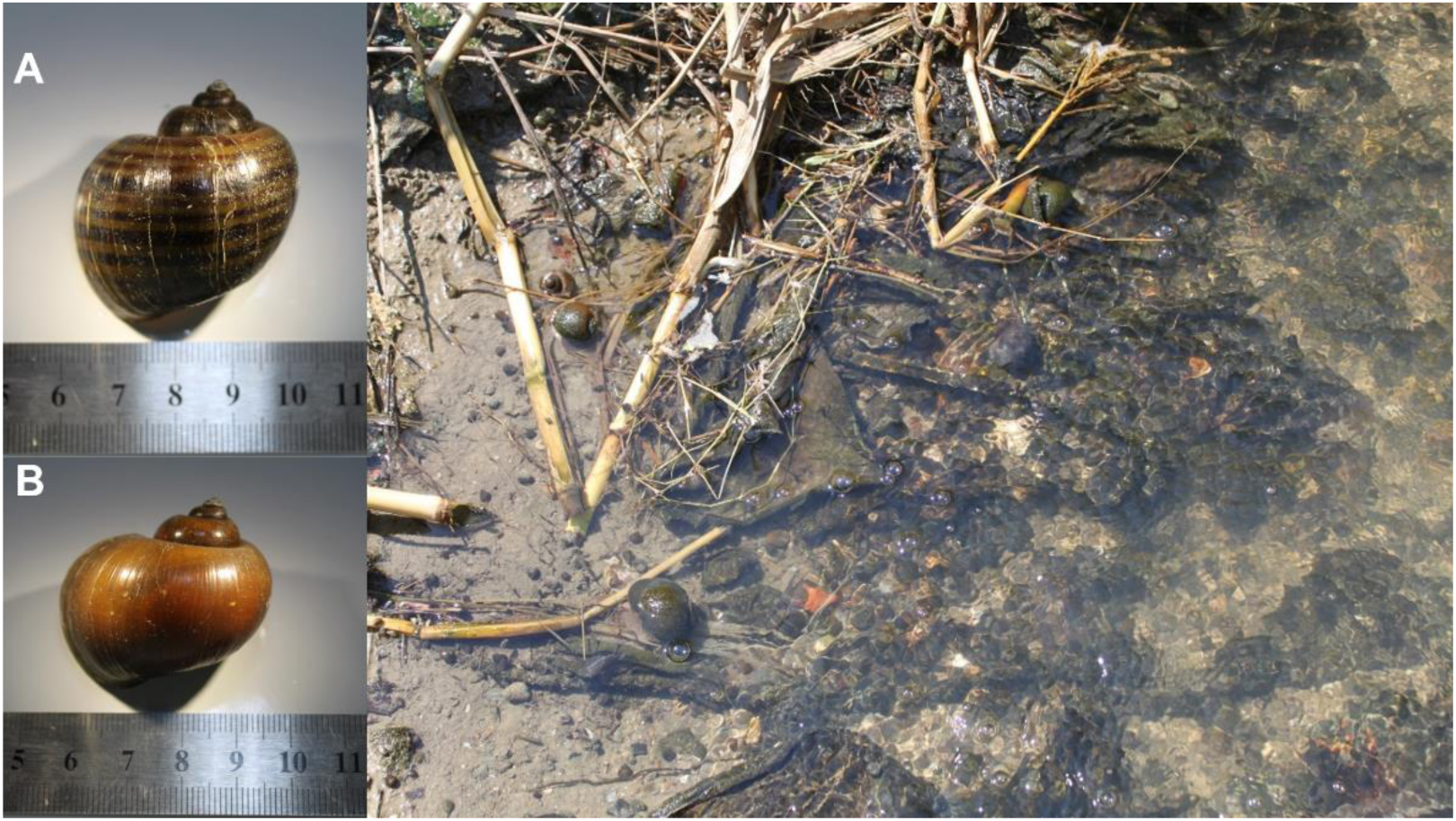
Classification of banded-brown apple snail (A) and yellow snail (B) based on shell color, and their activity in ditch habitats.

### 2.3. Statistical Analysis

Variance analysis of population density and individual phenotypes were performed after homogeneity of variance tests. Using the ArcGIS software (version 10.0, ESRI Inc., Redlands, CA, USA), the sampling sites were first located on an existing geographic information system (GIS). Subsequently, using the inverse distance-weighted method, a statistical technique for spatial prediction, a smoothed distribution map of *P. canaliculata* was produced based on the sample population density [26]. Using IBM SPSS software (version 20.0, IBM SPSS Inc., Chicago, IL, USA), correlations were tested among the five variables of density, body mass, shell height, sex ratio and color ratio. Spatial autocorrelation analysis was implemented with ArcGIS software (version 10.0, ESRI Inc., Redlands, CA, USA); Morans *I*, *z*-scores and *p*-values were recorded and comprehensively analyzed.

### 2.4. Ethics Statement

This study was carried out in accordance with the laws of wildlife protection, guidelines of investigation on aquatic animals, and experiment management policies in China. The observation, sampling and measurement were performed under the Certificate of Laboratory Animal Ethics approved by the Laboratory Animal Ethics Committee, Pearl River Fisheries Research Institute, CAFS.

## 3. Results

### 3.1. Spatial distribution, population density and population dynamics

The snail *P. canaliculata* had established natural population in all of the investigated sites, including the mountainous areas. Aquatic snail *Cipangopaludina cathayensis* (Heude, 1890), *Bellamya sp*., *Radix auricularia* (Linnaeus, 1758) and *Oncomelania hupensis* Gredler, 1881 were common accompanying species. As a lentic species, the snails prefer still waters to flowing streams. As a result, their abundance was notably higher in ditches, with a mean population density of 20.31 ± 11.55/m², compared to rivers and streams, where only sporadic discoveries of large individual snails were observed. There were clear differences in population densities among the examined sites, with the highest density of 49.00 ± 22.95 snails/m^2^ in Chaozhou and the lowest density of 4.14 ± 7.48 snails/m^2^ in Shantou (Figure 3). The population density was relatively low in Pearl River Delta with 8.22 snails/m^2^ in Guangzhou, 9.93 snails/m^2^ in Zhongshan, 8.83 snails/m^2^ in Jiangmen and 7.25 snails/m^2^ in Zhuhai, respectively. Observations revealed significant fluctuations in population density at the same location during extreme environmental conditions. For example, drought conditions led to the mortality of all snails, while floods tended to flush away the entire snail population.

**Figure 3.**
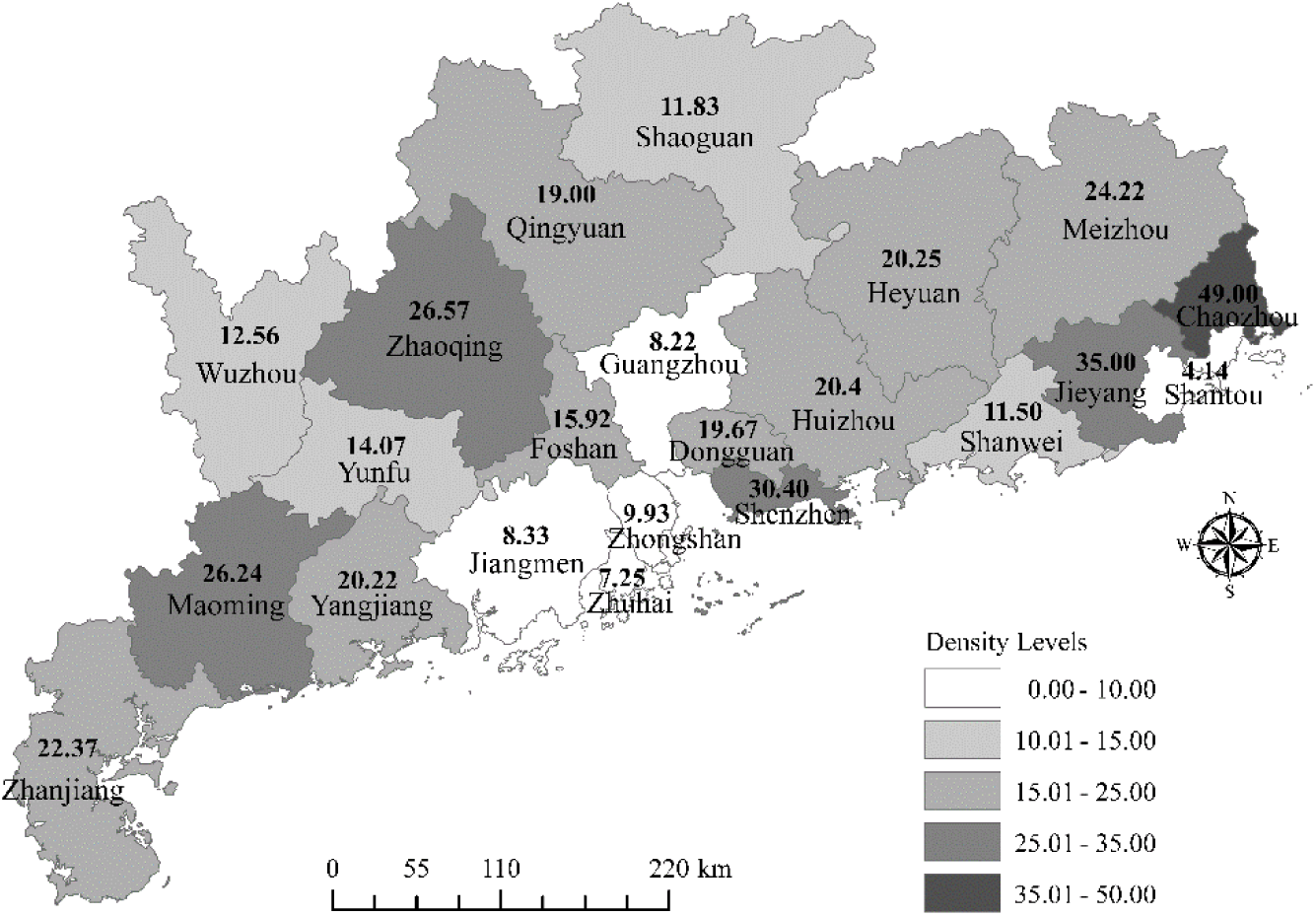
Spatial distribution of population density (snail/m²) for *Pomacea canaliculata*. Population abundance was categorized into five levels using the smart quantiles technique during the map creation process.

### 3.2. Geographic phenotypes

The main phenotypes of body mass, shell height, sex ratio and shell color ratio of the snail populations from the investigated ditches are shown in Figure 4 with averages shown at city level. The average weight of the sample snails (shell height >1.5 cm) was 8.93 ± 3.95 g with a range from the lightest 3.46 ± 0.10 g in Qingyuan to the heaviest 17.69 ± 4.35 g in Zhanjiang (Figure 4A). The overall mean shell height of the examined snails was 3.38 ± 0.66 cm with a range from 2.50 ± 0.14 cm in Qingyuan to 4.93 ± 0.46 cm in Foshan (Figure 4B). There was a clear female-biased sex ratio in the snail populations (overall mean female/male = 2.39 ± 1.01). The ratio varied from 3.94 ± 2.95 in Foshan to 0.93 ± 1.04 in Heyuan (Figure 4C). Shell color was classified as either banded-brown or yellow. Banded-brown snails were much more abundant than those with yellow shell color, with an average ratio of brown/yellow of 9.34 ± 7.52 (Figure 4D). Among the cities examined, Wuzhou and Yunfu were the only two where yellow snails were more abundant (brown/yellow ratios of 0.02 ± 0.01 and 0.22 ± 0.46, respectively). By contrast, the great majority of snails in Zhongshan were banded with a high ratio of 25.05 ± 13.44.

**Figure 4.**
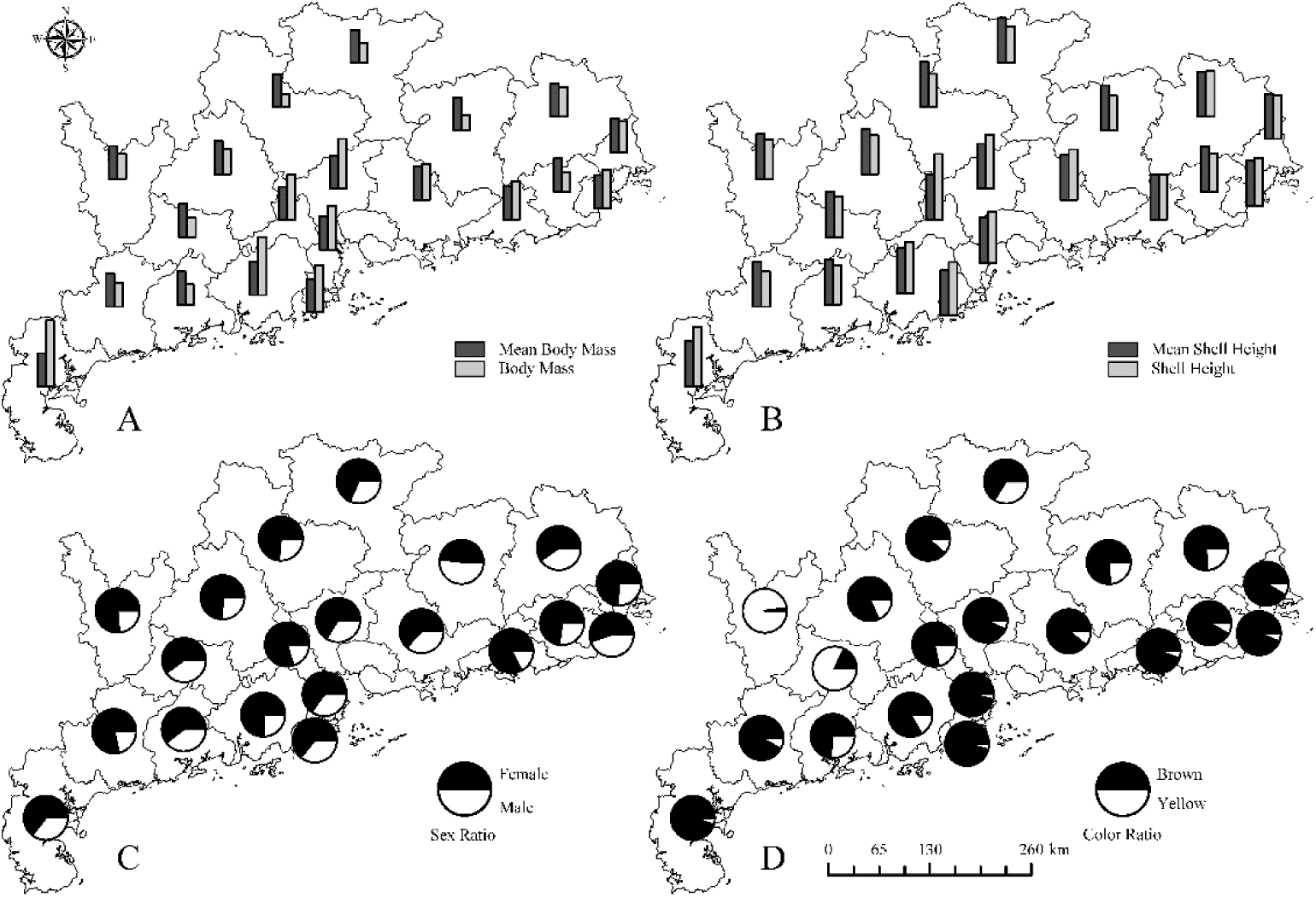
Phenotypic distribution analysis of *Pomacea canaliculata* across various sites, including body mass (A), shell height (B), sex ratio (C), and color ratio (D). The mean body mass and shell height, representing the average values across all investigated snails, serve as controls. Sex ratio is defined as the ratio of females to males, and shell color ratio is expressed as the proportion of brown to yellow shells.

### 3.3. Predictive analysis of the spatial variation of the P. canaliculata populations

Based on spatial analysis, a predicted map shows that the distribution and population abundance at the time of sampling was nonlinear and multi-centered (Figure 5). Clearly, Zhongshan, the initial site of introduction, was no longer the main output source of invasive snail populations. There was an area of low density extending from Shaoguan in the north to the Pearl River Delta in the south. The snail had established separate high-density populations in the eastern and western regions. A remarkably high-density region appeared in the Chaoshan plain in the east at a distance of 300 km from Zhongshan. Spatial autocorrelation analysis showed that the snails distributed nearly randomly and the population density was uncorrelated with a very low Moran’s Index of 0.059 (Table 1).

**Figure 5.**
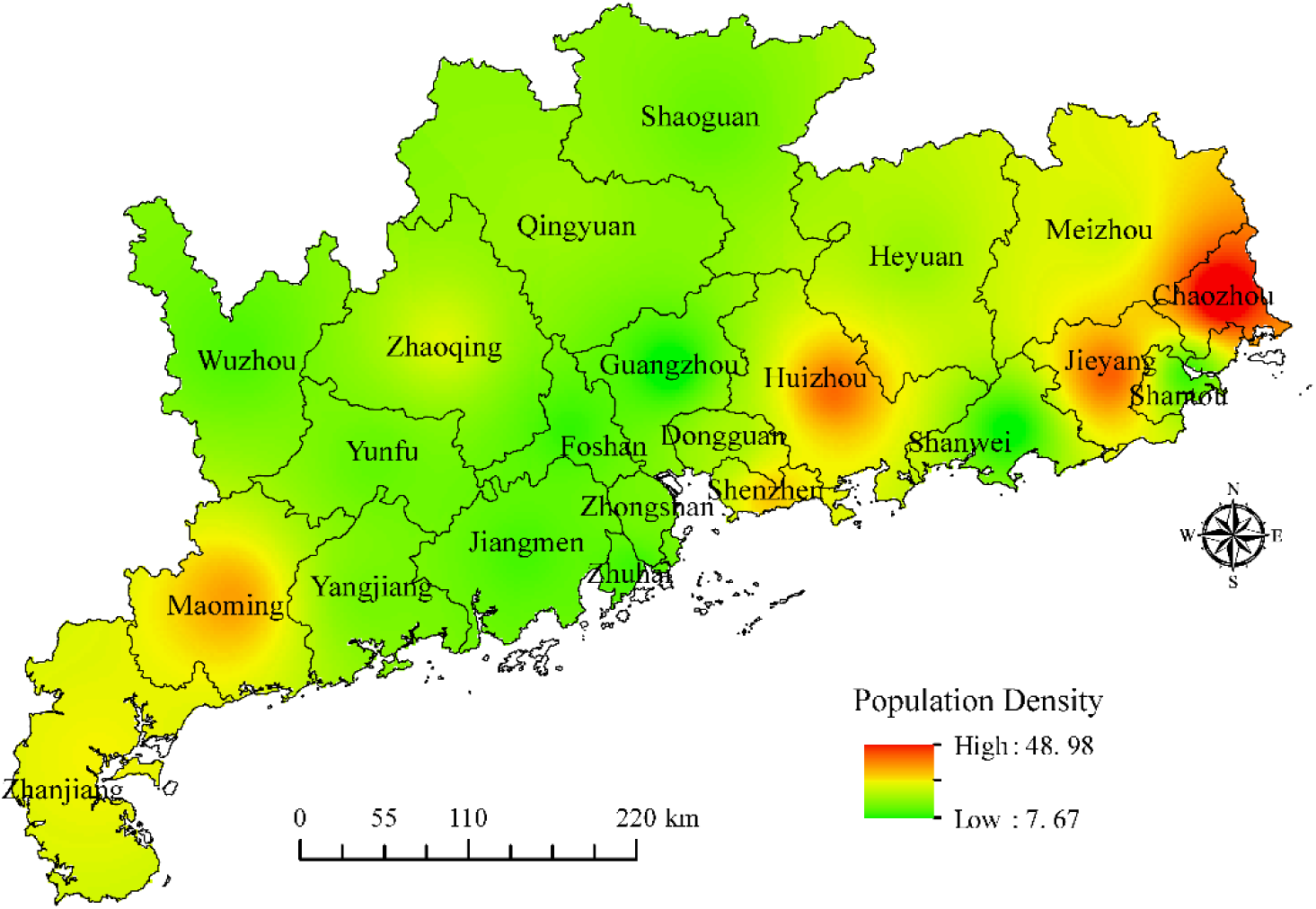
Modeled distribution of population density for *Pomacea canaliculata* in the investigated areas. The map was generated using the inverse distance-weighted method, with green representing the lowest density and red indicating the highest density.

**Table 1.** Spatial autocorrelation analysis of the population density and phenotypes of the snail *Pomacea canaliculata*. b/y = banded-brown/yellow, f/m = female/male.

| Variations | Moran's Index | z-score | p-value |
| --- | --- | --- | --- |
| Population density ( /m<sup>2</sup>) | 0.059 | 0.849 | 0.396 |
| Body mass (g) | 0.208 | 1.772 | 0.076 |
| Shell height (cm) | 0.268* | 2.196* | 0.028* |
| Sex ratio (f/m) | -0.296 | -1.670 | 0.095 |
| Shell color ratio (b/y) | 0.219 | 1.841 | 0.066 |

\* Significantly spatially autocorrelated.

Morphologically, the body mass and shell height of the snails in Pearl River Delta including Zhongshan in the middle of the study area and Zhanjiang in the west were distinguished from other parts (Figure 6A,B). Snails in these warm lowland areas were bigger than snails in inland areas. In contrast, the sex ratio changed discretely (Figure 6C). The distribution was multi-centered and discontinuous. With respect to shell color, snails with banded-brown shells occurred more frequently in coastal areas and yellow snails were predominantly distributed in inland mountainous areas (Figure 6D).

**Figure 6.**
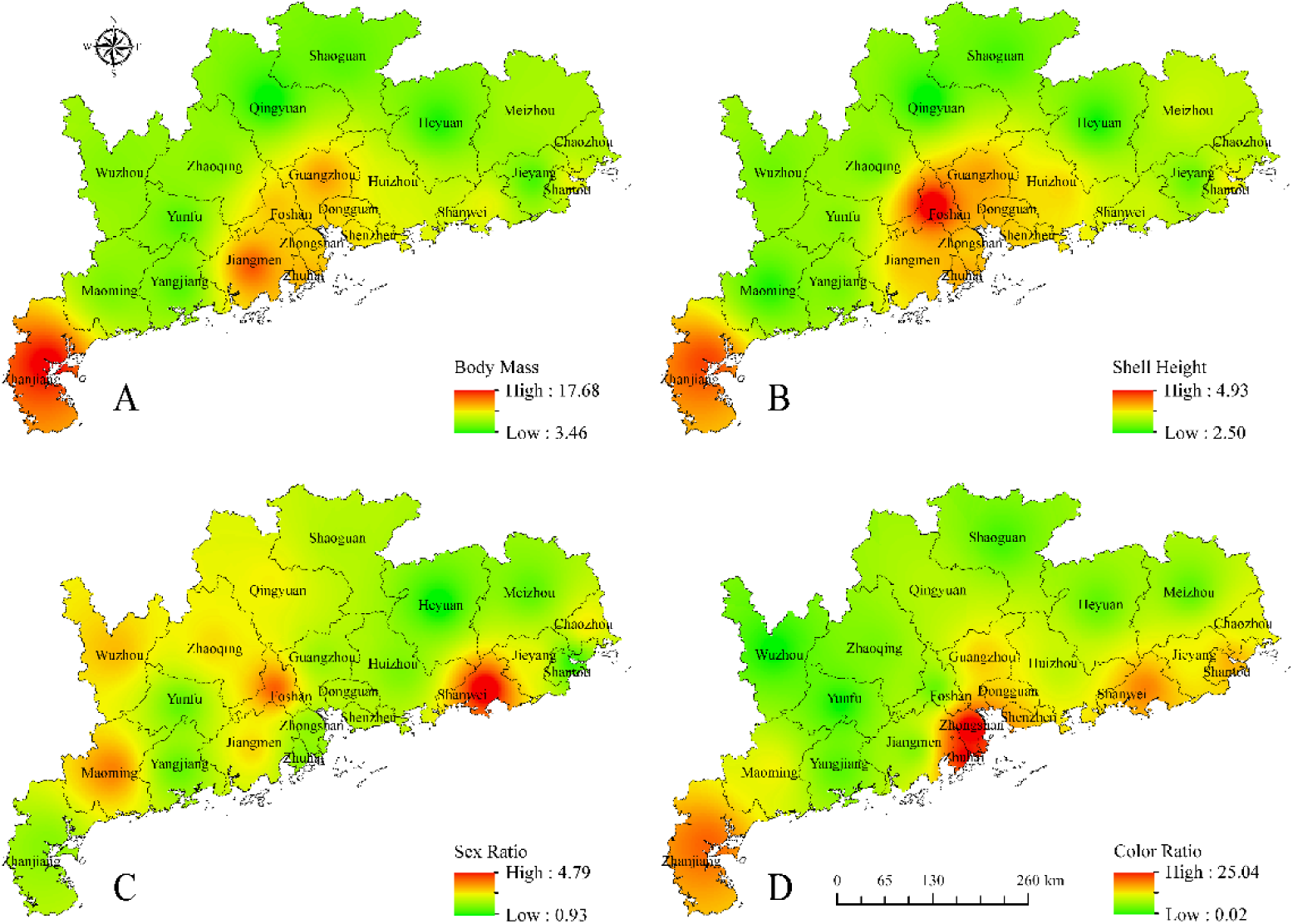
Predicted distribution of *Pomacea canaliculata* phenotypes in the study areas. (A) body mass (g); (B) shell height (cm); (C) sex ratio (female/male); (D) color ratio (banded-brown/yellow). The map was smoothed using the inverse distance-weighted method, with green indicating the lowest density and red indicating the highest density.

Spatial analysis of the phenotypic variations showed that only the autocorrelation of the shell height was statistically significant (Table 1). Body mass and brown/yellow-shell color ratio also indicated some spatial autocorrelation but this was not statistically significant. There was a negative autocorrelation in the sex ratio of the snail populations, but the z-score and *p*-value failed to reject the null hypothesis at a confidence level of 95%. Bivariate correlation analysis demonstrated that the body mass was significantly correlated with shell height (*r* = 0.88, *p* < 0.01) and with color ratio (*r* = 0.55, *p* < 0.05). However, no other significant correlations were found among the detected variables.

## 4. Discussion

Currently, many infectious diseases appear to be increasing, including vector-borne and water-borne diseases. Fluctuations in the dynamics of vector abundance are important determinants of vector-borne disease transmission [29]. This study showed that there were broad variations in the geographical distribution and phenotypes of populations of *P. canaliculata,* the main vector of *A. cantonensis*. These observations are consistent with a previous survey showing that the zoonotic nematode was widely distributed and that infection rates varied widely, from zero to 26.6% [18]. This ecological complexity must be largely determined by the snail bionomics. Furthermore, the temporal and spatial dynamics of vector snails interact with other host populations to enable parasite transmission [30]. Due to the complexity of angiostrongylosis epidemiology, this investigation into host ecology has the potential to enhance surveillance methods, bolster disease risk assessments, and contribute to more effective disease management strategies [31].

Although there was insufficient evidence to demonstrate that the invasion of *P. canaliculata* harmed the diversity of other snail species, we found that the local mollusks were always distributed accompanying with the alien apple snails in the environment. Therefore, there might be interactions among different mollusk species in a long term. *P. canaliculata* appears to be a more compatible host for *A. cantonensis* than the native snails *Cipangopaludina chinensis* (Gray, 1834) and *Bellamya aeruginosa* (Reeve) in which the infection rates were lower [18,32]. As much as 87 species of freshwater snails were recorded as the intermediate host snails of *A. cantonensis*. In China, 11 species of freshwater snails have been reported [33]. The coexistence and interaction between invasive snails and native species could be important biotic factors affecting the transmission of the infectious disease. Additionally, high densities of the apple snails were associated with a complete shift in both ecosystem state and function [34]. The detrimental impact of *P. canaliculata* on ecosystems suggests that the invasion of the snail may promote the transmission of *A. cantonensis* through its effects on biodiversity [35].

Limiting factors for the distribution of *P. canaliculata* are considered to be mainly chemical and physical [9]. *P. canaliculata* is adapted to all types of freshwater bodies, except for large rivers. Previously, it was shown that there are substantial differences in chemical and physical parameters among streams, ditches, ponds, paddy fields and marshes [9]. Snails mainly accumulate in and spread through ditches, which are their most common habitats. Therefore, as their population density and abundance are highest in ditches, the present survey focused on ditches. However, it is worth noting that some sites in abandoned wet rice paddies also contained a high density of medium-sized snails, suggesting that there is a high risk of them spreading into unexpected situations.

Environmental factors greatly affected species population dynamics, leading to large differences in distribution patterns [36]. The geographical ranges of many species, including many infectious disease vectors and intermediate hosts, are assumed to be constrained by climatic tolerances [37,38]. The variation in the distribution of *Pomacea sp*. could be attributed to differences in cold tolerance [39]. However, it is noteworthy that all the surveyed sites shared tropical or subtropical monsoon climates characterized by long summers and abundant rainfall. This aligns with the observed widespread distribution of apple snails across all investigation sites. However, as a lentic species [40], the population density would be habitat-determined during warmer times, related to the abundance and availability of food resources. Additionally, growth and reproduction of the apple snail is density-dependent, while grazing is size-dependent [41,42]. Therefore, its preferences for abundant macrophytes and periphyton, the depth of water, adhesion characteristics and its behavioral adaptation to these conditions may all contribute to the marked population density differences among investigated sites [43].

Our data revealed a geographic stratification of the phenotypic variations of *P. canaliculata* of body mass, shell height and color ratio. It has been reported previously that spatiotemporal morphological variations are frequently present within the same species of aquatic gastropods [44]. The distribution, abundance, and morphology of *Pomacea sp*. are significantly influenced by water temperature and habitat characteristics. Consequently, variations in shell height and weight are likely attributed to ecophenotypes related to temperature fluctuations [45, 46]. Populations in regions such as Chaoshan Plain, the Pearl River Delta Plain, and coastal areas experienced lower environmental pressure from temperature. This resulted in a notable spatial autocorrelation of shell height variation, indicating that snails in these areas tended to have longer lifespans and achieve larger sizes, particularly in lower-altitude regions. Nevertheless, phenotypic traits are conservative [47]. Even *A. cantonensis* larvae from young snails could also infect rats successfully. The worm burden differed with snail body size [20]. More research on the potential role of phenotypic variation in angiostrongyliasis transmission may improve the risk assessment of human eosinophilic meningitis. Another notable finding was the difference in shell color varying from yellow to banded-brown. The yellow-color snails were more distributed in the inland mountainous areas, whereas banded-brown snails were mainly located in the downstream areas. There may be some intense biological and genetic interactions among the groups in the intermediate regions. Previous research showed that the yellow phenotype follows simple Mendelian inheritance, being recessive to brown [48]. Though, there was no scientific reports. A deliberate introduction of yellow-snail for aquaculture in the 1980s probably explains the original specific color distribution. However, more phylogenetic evidence is heavily needed to examine the origin of different colored snail populations. Taking snail color into consideration may improve the assessment of susceptibility of *P. canaliculata* to *A. cantonensis* [19]. Partly because we chose to investigate shell heights ≥ 2.5 cm, and ditches as the habitat, the total sex ratios of the populations were higher than in other reports from paddy fields [42]. Additionally, it has been demonstrated experimentally that the sex determination in the apple snail *P. canaliculata* was oligogenic [49]. Female-biased sex ratio was found to be significantly positively associated with temperature and precipitation at the region level of 14 cities in Guangdong province [50]. In this investigation, we observed that the sex ratios of *P. canaliculata* were highly variable among small populations and that this variation was not geographically correlated. Hence, it could be inferred that the snails had already established small populations exhibiting a degree of genetic differentiation. This was consistent with the mating experiment that the sex ratios bias towards females showed greater in the inbred lines than in the outcrossed lines [51].

The current investigation complements an earlier study of the spatial epidemiology of *Pomacea*-related zoonotic parasitic diseases [7,16]. It may help to control outbreaks of snails by monitoring their populations to devise a strategy for managing invasive snails for public health and economic safety. Although there was insufficient evidence to show that a high snail population density indicated a higher prevalence of *A. cantonensis* infection in humans, almost all known cases have been associated with consumption of raw snails. As an emerging foodborne parasitosis, it is necessary to monitor populations, to track their possible dispersal routes, and to study the characteristics of *A. cantonensis* infection among different geo-ecological groups in the upcoming studies. Transmission of vector-borne diseases is driven by a multitude of epidemiological, ecological and socioeconomic factors. Future risk assessment should take into account the complexity of transmission dynamics in vector-pathogen systems [52]. Further ideas for controlling snails could be considered. Specifically, it is important to analyze the annual dynamics of infection rates and to study the susceptibility and compatibility of infection of *P. canaliculata* with *A. cantonensis* based on shell color, longevity, habitat and gender. In addition, there should be more research into the effects on infection of key physicochemical and environmental factors, such as pH, temperature and precipitation. Special attention should be directed toward snail control and management in mountainous areas [52]. Additionally, forecasting the future distribution, dynamics and dispersal capacity of the vector snails under population evolution, land-use change and climate change is necessary to concerning the control of this emerging infectious disease [53].

## 5. Conclusions

In summary, the current study is the first to document the distribution pattern of *P. canaliculata*, the invasive vector of *A. cantonensis*, around its initial introduction site in China, its population abundance and related phenotypic variation. We show that, 30 years after their introduction, the snails were already stably established at all of the investigated sites, even in mountainous areas. The mean population density was 20.31 ± 11.55/m^2^ ranging from the highest 49.00 ± 22.95 snails/m^2^ to the lowest 4.14 ± 7.48 snails/m^2^. The distribution of population abundance was nonlinear and multi-centered. The average shell color ratio of brown/yellow was 9.34 ± 7.52 ranging from 0.02 ± 0.01 to 25.05 ± 13.44. There was a geographic stratification for the phenotypic variations of *P. canaliculata* in body mass, shell height and shell color ratio. Bivariate correlation analysis demonstrated that the color ratio was significantly correlated with body mass (*r* = 0.55, *p* < 0.05). All of the surveyed parameters were highly variable, which implies strong adaptability and phenotypic plasticity of the snail, and emphasizes the complexity of the parasite transmission system. Geographic variations in the distribution, abundance, habitat preference, and key phenotypes of the snail may have implications for devising cost-effective control interventions for the infectious *A. cantonensis*. Snail control interventions should take into account the spatial distribution of snail densities and the temporal and spatial variations of abundance and phenotypes.

## Author Contributions

Conceptualization, D.L., Y.H., and J.L.; methodology, D.L. and H.Z.; software, D.L. and H.Z.; validation, D.L. and H.Z.; formal analysis, D.L. and H.Z.; investigation, D.L., H.Z., D.G., X.M., H.S., Y.Y. and M.X.; writing—original draft preparation, D.L. and H.Z.; writing—review and editing, D.L.; visualization, D.L. and H.Z.; project administration, D.L. and Y.H.; funding acquisition, D.L. and Y.H. All authors have read and agreed to the published version of the manuscript.

## Institutional Review Board Statement

This study was carried out in accordance with the laws of wildlife protection and guidelines of investigation on aquatic animals in China. The animal study protocol was approved by the Laboratory Animal Ethics Committee of Pearl River Fisheries Research Institute, CAFS (LAEC-PRFRI-2017-01-01).

## Informed Consent Statement

Not applicable.

## Data Availability Statement

Data are available on request.

